# Quantitative basis of meiotic chromosome synapsis analyzed by electron tomography

**DOI:** 10.1101/705764

**Authors:** Marie-Christin Spindler, Sebastian Filbeck, Christian Stigloher, Ricardo Benavente

## Abstract

The synaptonemal complex is a multiprotein complex, which mediates the synapsis and recombination between homologous chromosomes during meiosis. The complex is comprised of two lateral elements and a central element connected by perpendicular transverse filaments (TFs). A 3D model based on actual morphological data of the SC is missing. Here, we applied electron tomography (ET) and manual feature extraction to generate a quantitative 3D model of the murine SC. We quantified the length (90 nm) and width (2 nm) of the TFs. Interestingly, the 80 TFs/μm are distributed asymmetrically in the central region of the SC challenging available models of SC organization. Furthermore, our detailed 3D topological analysis does not support a bilayered organization of the central region as proposed earlier. Overall, our quantitative analysis is relevant to understand the functions and dynamics of the SC and provides the basis for analyzing multiprotein complexes in their morphological context using ET.

## Introduction

Meiosis is the special cell division that results in the generation of haploid cells, which are required for sexual reproduction. Haploidization is achieved during the first meiotic division. In the process, homologous chromosomes have to synapse and recombine before they can segregate properly. The synaptonemal complex (SC), a meiosis-specific and evolutionarily conserved structure of meiotic chromosomes hereby stabilizes the synapsis of homologous chromosomes and is further required for the progression of recombination.

The ladder-like structure of the SC comprises of two lateral elements (LEs) that are connected to each other via the central element (CE) and numerous perpendicularly running transverse filaments (TFs) [1–4]. The chromatin corresponding to homologous chromosomes is attached to LEs in regular arrays of DNA loops [5, 6].

The LEs -also called axial elements prior to chromosome synapsis- are the first components of the SC to assemble early during meiotic prophase I [6, 7]. In mice, the major protein components of the LEs are SYCP2 (1505 amino acids) and SYCP3 (257 amino acids) [8, 9]. Crystallographic analysis of SYCP3 showed an elongated tetrameric assembly of antiparallel α-helices of 20 nm length. This antiparallel configuration leads to the exposure of the N-terminal DNA binding sites at both sides of the SYCP3 protein. [10]. Furthermore, it has been demonstrated that SYCP3 self-assembles into regular higher-order superlattices [11–13] and that the tetramer is able to compact chromatin by stabilizing DNA loops in vitro [10]. SYCP2 also contains multiple potential DNA binding motifs, yet very little is known about its role in meiotic chromosome organization [8, 14, 15]. SYCP2 interacts with both SYCP3 and SYCP1, the main component of TFs [16, 17, 11, 18]. SYCP3 and SYCP1, however, do not interact directly with each other [16, 17, 11]. Therefore, by binding to both SYCP1 and SYCP3, SYCP2 likely provides sites for TF binding to the LE [18].

In mammals, TFs are formed by homodimers of SYCP1 (976 amino acids) [19] with SYCP1 molecules showing the same polarity. The C-termini of the SYCP1 molecules interact with components of the LE [16, 20] and contain basic patches separated by approx. 30 Å, indicating DNA binding properties [21]. Truncated SYCP1 dimers, lacking this region show no interaction with DNA [21]. The two long α-helices of the SYCP1 dimer, which make up the bulk of the protein, form a coiled-coil structure (amino acids 101-783) connecting either one of the LEs with the CE [16, 20]. Crystallographic data recently revealed a length of the α-helical SYCP1 core of 900 Å [21]. Polymerization studies of SYCP1 are consistent with the notion that the length of the coiled-coil is a primary determinant of SC width [22]. In the CE, TFs attaching to opposite LEs interact with each other via the non-helical N-terminal ends of SYCP1 dimers [16, 20, 21].

SYCP1 also plays an important role during SC assembly as it essential for recruiting the different CE protein components [23]. Previous studies concluded that SYCP1 first recruits CE proteins SYCE3 and SYCE1 and then SYCE2-Tex12 in sequential order [24, 7]. Interaction studies between the N-terminus of SYCP1 and SYCE3 support this model of SC assembly [25, 26, 27]. Independent assays confirmed the assignment of subsequent CE interacting partners to the two subsets SYCE3-SYCE1 and SYCE2-Tex12. SYCE1 e.g. interacts with another component of the CE, SIX6OS1. [24, 28, 29, 27, 30]. Attempts to analyze distributions of these proteins within the CE using immunogold EM suggested a bilayered organization of the SC central region [31].

In the last more than 60 years, a variety of studies has been conducted to elucidate the molecular organization of the SC. The early approaches were mostly based on electron microscopy (EM) [3]. They were complemented by immunolocalizations, providing valuable insight into the molecules’ identity and localization of SC proteins [32]. Pushing the resolution of light microscopy beyond the limit of diffraction, techniques such as *d*STORM enabled precise protein localization maps with nanometer resolution [33, 34]. However, these super-resolution approaches lack the morphological context that EM techniques can provide. EM tomography (ET) bridges the gap between nanometer resolution and morphology. ET allows the resolution of the macromolecular architecture of SC components in both the *xy*-axis and the *z*-axis, as previously shown by Schmekel et al [35, 36]. However, at that time it was technically impossible to perform direct 3D reconstructions of the recorded structures. Furthermore, it was not possible to annotate the structures within the SC or to provide detailed quantitative data. Nowadays, improved hard- and software enables computerized reconstruction, annotation as well as feature extraction and quantification. Here, we have taken advantage of technical progress in order to provide a detailed structural and quantitative analysis of the SC. These data are of relevance in order to understand the functions and dynamics of the SC.

## Results

As mentioned above, in the mouse, many of the SC protein components are already known and detailed protein localization maps with nanometer precision have been generated [33] (Figure 1). However, our knowledge about the quantitative aspects of the SC organization is rather limited [4]. To this end, we have investigated the 3D ultrastructure of the murine SC by using electron tomography (ET). Dual-axis electron tomograms were acquired, reconstructed and manually segmented to create 3D models (Figure 2; S1/S2 Movie). The point data extracted from these models were processed through custom Matlab scripts for quantification of TF features (see Methods) [37]. The quantitative data presented here correspond to six SC tomograms and the 501 TFs contained therein. Individual tomograms are comparable with each other as they correspond to the early pachytene stage of 14 days-old mice.

**Figure 1.**
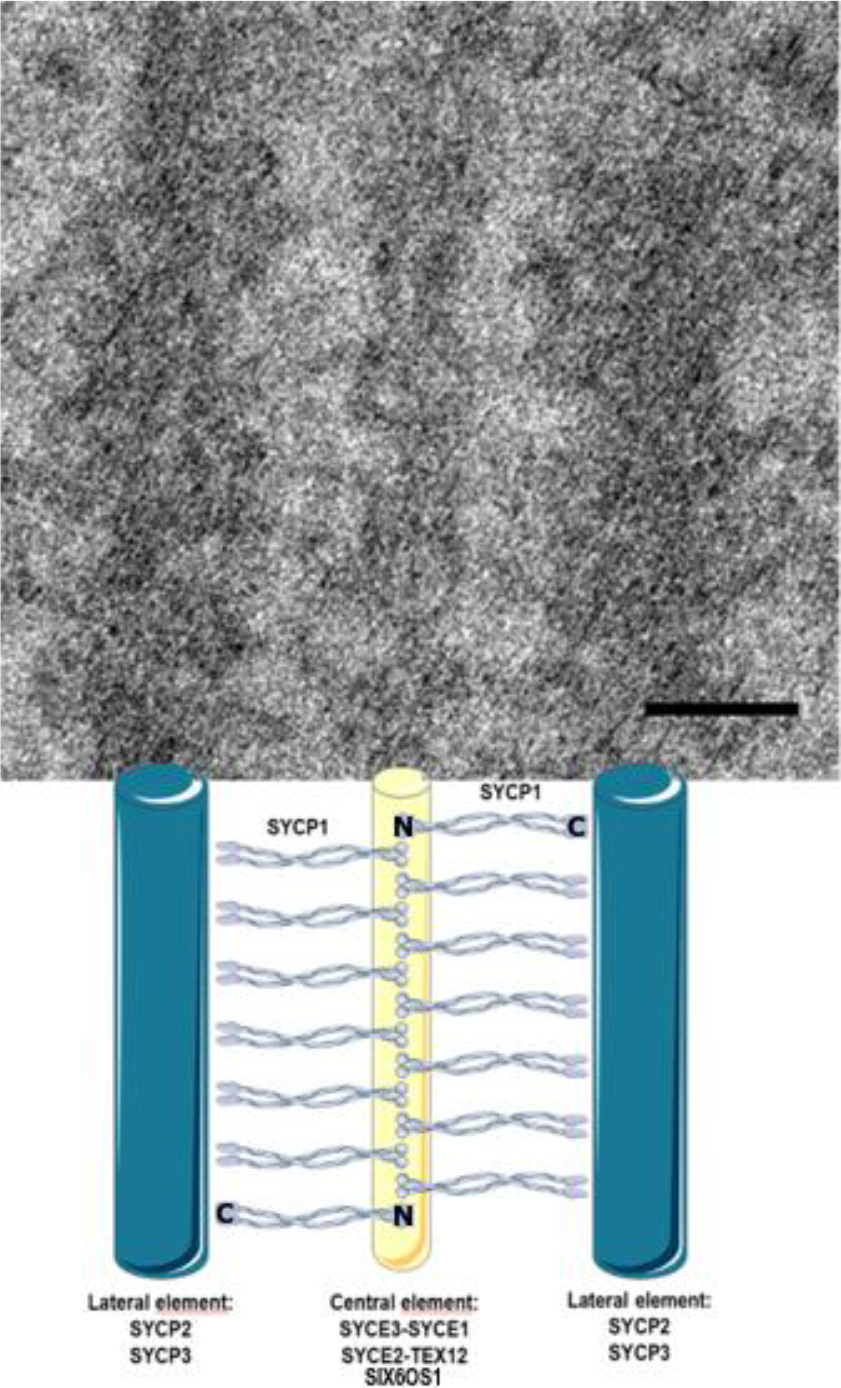
Synaptonemal complex. Electron micrograph of the synaptonemal complex combined with a schematic representation of its protein components: two lateral elements comprised of SYCP2 and SYCP3 in dark blue, central element (SYCE3, SYCE1, SIX6OS1, SYCE2, and TEX12) in yellow. The main component of the transverse filaments, SYCP1 (light blue), is represented as a coiled-coil with the N-terminus of the protein residing in the central element and the C-terminus in the lateral element.

**Figure 2.**
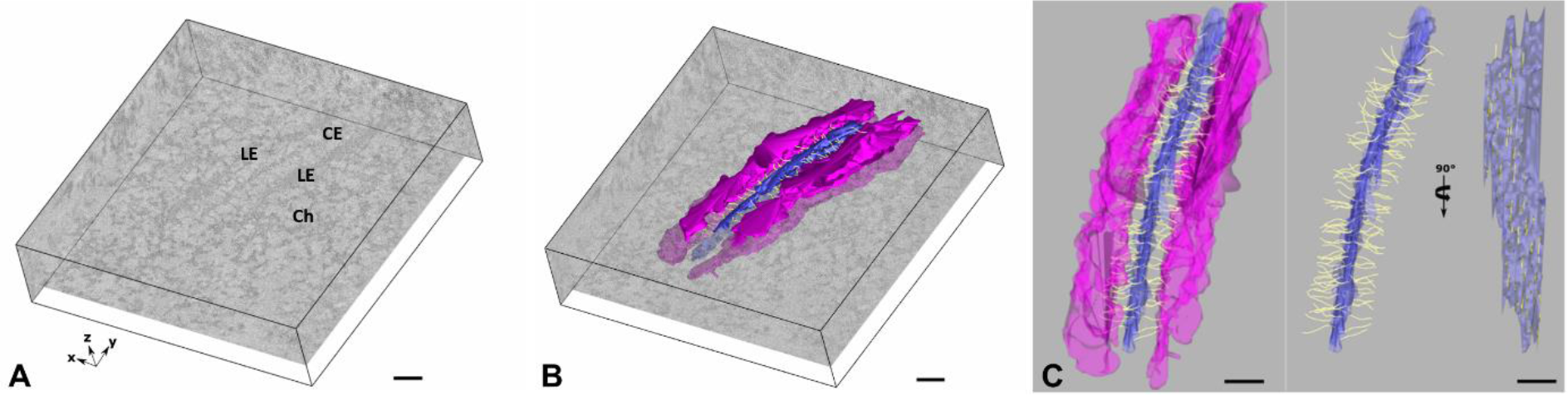
Generation of EM-derived 3D model of the synaptonemal complex. Scale bar: 100 nm. A. Virtual section of the reconstructed electron microscope tomogram containing the tripartite synaptonemal comple
x. LE: lateral element, CE: central element, Ch: Chromatin. Coordinate system representation indicating the three-dimensional orientation of the tomogram. B. 3D model of the synaptonemal complex generated by manual generation embedded in a virtual section of the tomogram. Magenta: lateral element; blue: central element; Transverse filaments in yellow. C. Left: Frontal view of the 3D model of the synaptonemal complex. Right: Frontal and lateral view of the synaptonemal complex without the lateral elements. Magenta: lateral element; blue: central element; Transverse filaments in yellow.

We first measured well-known components of the SC (Figure 3). According to our data, the width of the central region (i.e. from the inner edge of a LE to the inner edge of the LE of the opposite side of the SC) is 114 nm (± 17 nm). From the inner edge of a LE to the edge of the CE, the distance is 45 nm (± 10 nm). Correspondingly, the width of the CE is 29 (± 8 nm). Under our experimental conditions, the width of TFs is 1.9 nm (± 0.3 nm). These results are in accordance with previous transmission EM studies [1–3] and early EM tomographic data [4].

**Figure 3.**
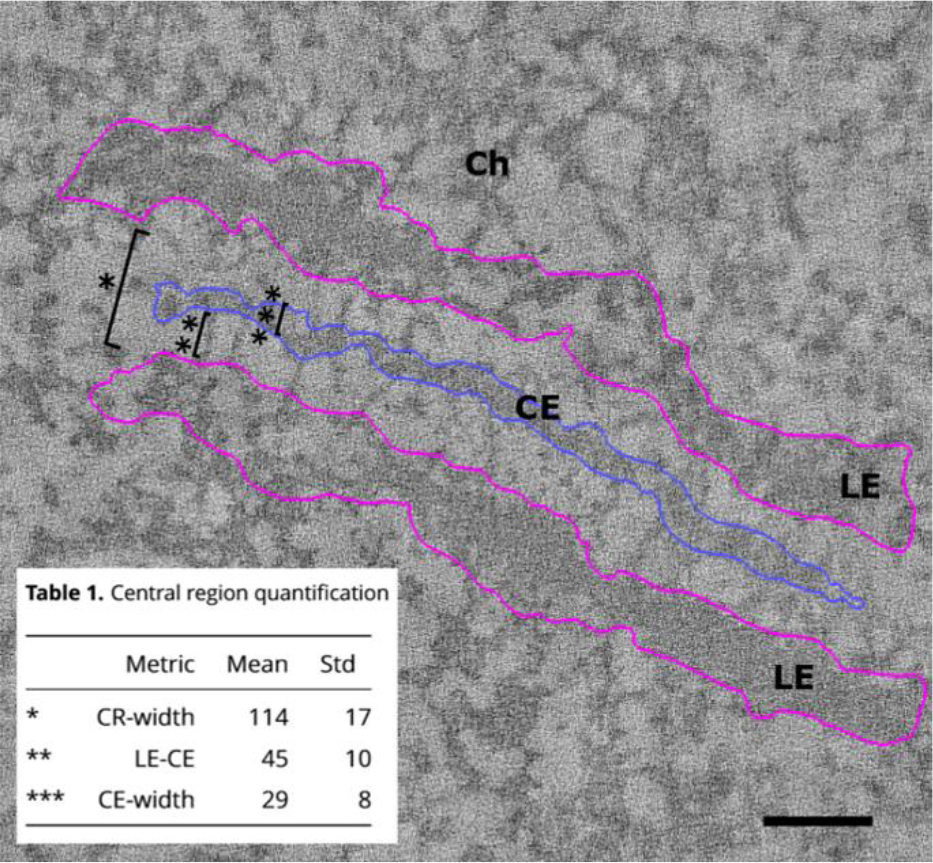
Quantification of synaptonemal complex dimensions. Visual representation of measured distances in nanometer within the synaptonemal complex. The width of the central region (*, CR-width), the distance between the lateral and central element (**, LE-CE) and the width of the central element (***, CE-width) were determined. Inset: Table 1 containing the central region quantification. LE: lateral element, CE: central element, Ch: Chromatin. Scale bar: 100 nm.

The 3D reconstructions of synaptonemal complexes enabled us to determine the number of TFs along a SC, i.e. 79 TFs/μm (± 8), resulting in a density (measured at the CE insertion site of TFs) of 1046 TFs/μm^2^ (± 253) (Figure 4). The number of TFs per micrometer in the mouse corresponds to the upper limit of the reported 50-80 TFs/μm of hamster synaptonemal complexes [38].

**Figure 4.**
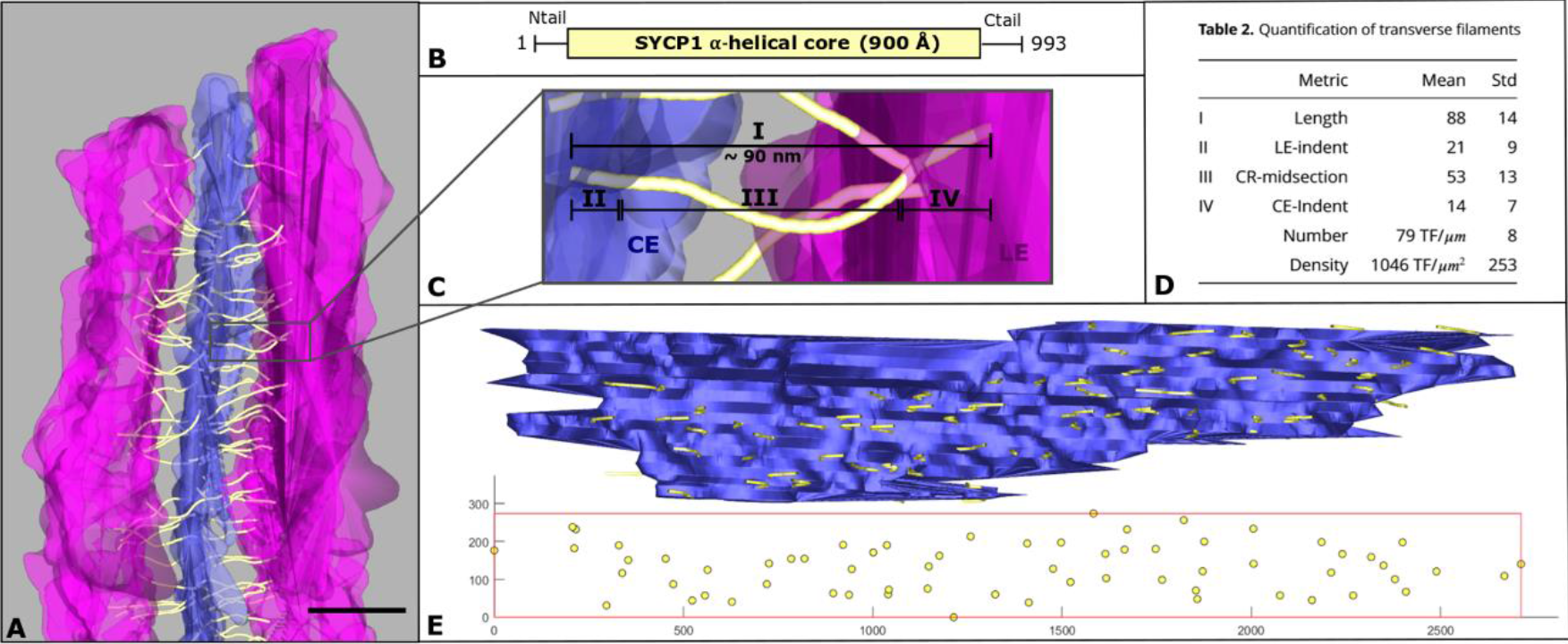
Quantification of transverse filaments. A. Frontal view of a section of the 3D model of the synaptonemal complex. The lateral element in magenta, the central element in blue, the transverse filaments in yellow. Scale bar: 100 nm. B. Sequence of transverse filament main component SYCP1. In mice, the protein has a length of 993 amino acids. The majority of SYCP1 makes up the alpha-helical core of the protein, which in turn has a size of 900 angstroms. The protein is flanked by two unstructured sequences at the N- and C-terminus. C. Magnified view of a single transverse filament of A. The average length of a transverse filament is roughly 90 nm (I). The filament is embedded both in the central element (CE, II) and in the lateral element (LE, IV), completed by a larger midsection in the central region (III). D. Table 2 Quantification of transverse filaments. Table contains the average length of the transverse filaments in nm (TF, I), the length of the part of the filament embedded in the lateral element in nm (LE-indent, II) and the central element in nm (CE-indent, IV), the length of the midsection of the TF in the central region (CR-midsection, III) as well as the number of filaments per μm and μm^2^ synaptonemal complex. E. Top: Central element and the transverse filaments embedded in one of the lateral elements shown in the lateral view. Bottom: 2D projection of the endpoints of the transverse filaments in the lateral element. A box is fitted through the four outer points to give an estimate of the transverse filaments per area of synaptonemal complex (in pixel).

Interestingly, the higher resolution in the z-axis provided here by ET enabled us, for the first time, to track the TFs also within the environment of the central and the lateral elements (Figure 4). Thus, the average length of a TF is 88 nm (±14 nm). Remarkably, this value is in accordance with crystallographic data of the coiled-coil domain of SYCP1, the protein component TFs. The respective study stated a length of the SYCP1 coiled-coiled region of 900 Å [21]. Furthermore, we were able to determine the length of the TF subsection within the environment of the CE (14 nm ±7 nm) and the LE (21 nm ±9 nm). In accordance with these data, the TF segment between a LE and the CE is 53 nm (±13 nm) long (Figure 4). Given that TFs consist of protein SYCP1 [22], our segmentation would indicate that the coiled-coil domain of a SYCP1 dimer reaches into the CE and the LE.

SCs are usually represented as symmetric ladder-like structures showing TFs of the one side interacting 1:1 in the CE with TFs of the other side [4]. Here, we were able to demonstrate that this is not necessarily the case. We counted the number of TFs in a SC segment and found differences from up to 21% when comparing the number of TFs between the “right side” and the “left side” of a SC. The consequence is that in the CE a certain TF of the “right side” may lack a partner in the “left side” (Figure 5, ‘single’ TF model). In other terms, a SYCP1 dimer would be able to integrate into the CE without the necessity of a SYCP1 dimer on the opposite side. Generally, we find three frequent arrangements of transverse filaments in the CE, i.e. TFs with a neighbor on the same side (‘parallel’), the opposing site (‘opposite’) and TFs without a partner (‘single’). Because of these findings suggesting asymmetries, we became also interested in knowing the minimal distance between TFs (Figure 5). To this end, we considered the CE-endpoint of a certain TF and measured the distance to the endpoint of the next opposing TF (i.e. originating from the opposite LE) as well as to the endpoint of the next parallel TF (i.e. originating from the same LE). The average minimal distance was of 17 nm (± 9 nm) for opposite TFs and 20 nm (± 10 nm) for parallel TFs. The observed difference between the two data sets is statistically significant (p = 4.63 × 10^−10^, Wilcoxon rank-sum test). We also measured the minimal distance of TF endpoints in the LEs. Here, the average minimum distance was 21 nm (± 10 nm). There was no significant difference in comparison to parallel TFs in the CE (p = 0.0438, Wilcoxon rank-sum test).

**Figure 5.**
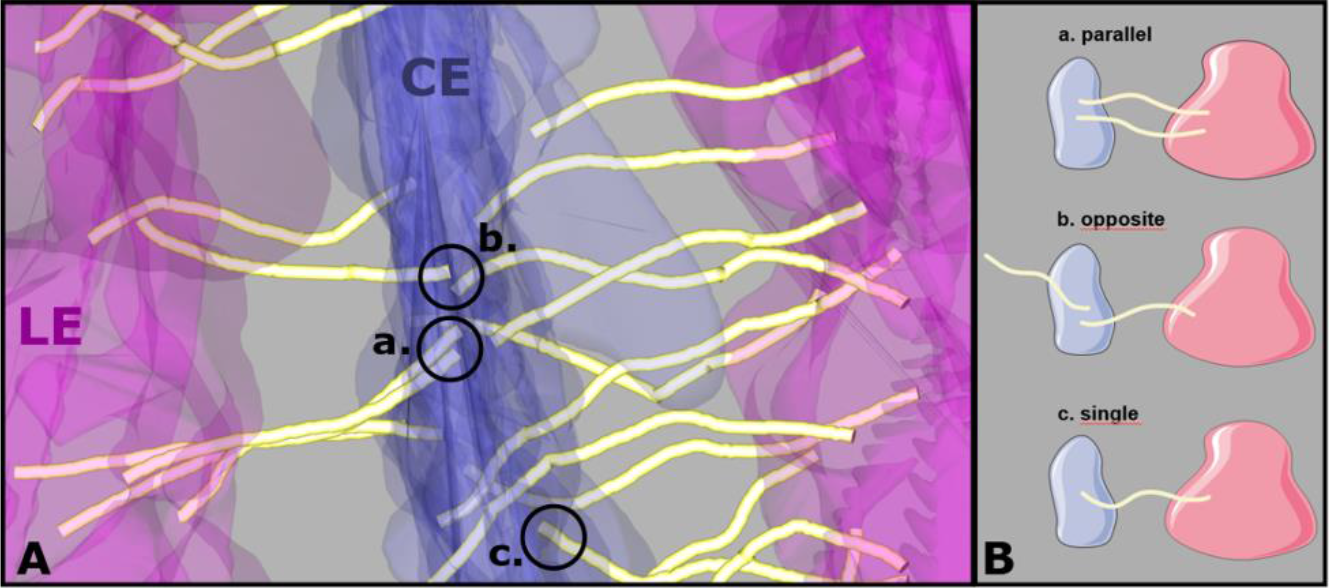
Transverse filament pairings in the central element. A. Section of the synaptonemal complex showing the three representations of transverse filament pairings in the central element: a. neighboring filaments originating from the same lateral element; b. neighboring filaments originating from different lateral elements; c. transverse filaments without a partner filament on either side. B. Model of transverse filament pairings in the central element derived from transverse filament nearest neighbor distributions in the central element: a. parallel, b. opposite and c. single.

ET in insects has revealed the separation of the central element into three to four distinct components in a cross-sectional view. The transverse filaments (TFs) pass through these components, which predetermines the organization of TFs into three to four layers [35, 36]. In mammals, the central element appears amorphous [3]. The lack of clearly separated central element components impedes the visual assignment of TFs into layers. In this case, a detailed topological analysis of the TFs requires the generation of a 3D model, which was technically impossible at the time of previous ET studies. However, immunoelectron and structured illumination microscopy (SIM) approaches in mouse showed bimodal distribution patterns of SYCP1 (the major TF component) in the central element that suggest a bilayered TF organization [31, 33].

The technological advances of recent times have allowed us to generate a 3D ET derived model of the synaptonemal complex in the mouse. Based on this model we are now able to address the question of TF distribution at ET level. Visual inspection of the TFs in the cross-sectional view did not reveal distinct layers of TFs in mouse (Figure 6A). To provide a more direct visualization of TF distribution, we proceeded to inspect the intersection points of the TFs with the CE in the lateral view. Due to the three-dimensional nature of the CE, the intersection points may reside in different z-planes, which can lead to misinterpretation due to superimposition. We, therefore, generated a 2D projection of the intersection points of TFs with the CE. Here, no distinct TF distribution pattern was observed (Figure 6B). Since visual inspection alone can be error-prone, we additionally approached the question for TF layers from a mathematical standpoint. Under the assumption that the TFs are organized into two layers in mammals, as suggested in previous studies, we fitted two planes through the endpoints of each TF. The two sets of filaments on either side of the central elements were hereby treated independently. In the presence of two layers of TFs, a parallel orientation at a designated spacing is expected. Instead, the fit supports the organization of the TFs into a single layer for all six analyzed tomograms. The exemplary fit displayed in Figure 6C assigned the TFs to two layers along the course of the SC through the section, which almost forms a single plane, instead of the expected two parallel planes. To conclude, neither through visual inspection of the TFs in the cross-sectional view nor through inspection of the 2D projection of intersection points of TFs with the CE or by the fitting of 3D planes we were able to detect any distinct layers of TFs in the mouse.

**Figure 6.**
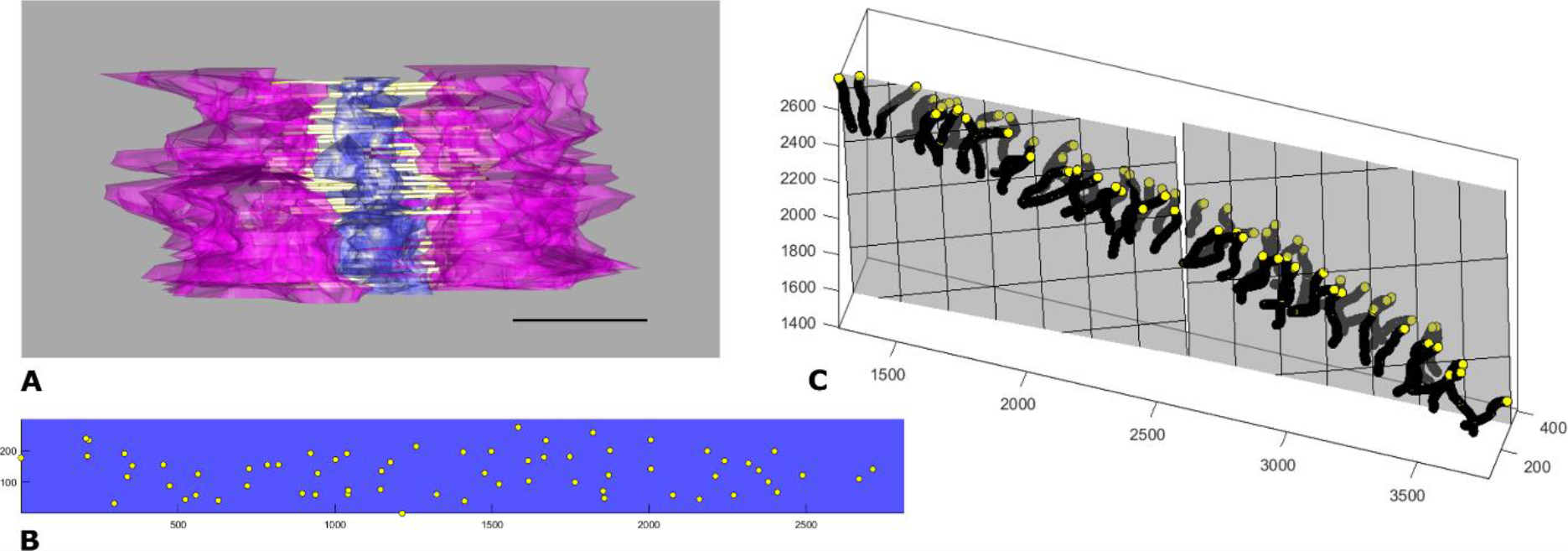
Mathematical modeling of EMT derived 3D model refuses multilayered transverse filaments of the synaptonemal complex. A. Cross-sectional view of the synaptonemal complex with no apparent layered organization of the transverse filaments (TFs). Magenta: lateral elements, purple: central element, yellow: TFs. Scale bar: 100 nm. B. 2D projection of the intersection points of the transverse filaments (TFs, yellow) with the central element. The area of the central element containing TFs shown in purple. In this view, no layers of TFs are apparent. C. Fit of two planes to the endpoints of the transverse filaments (TFs). A layered organization of the TFs would be reflected in two parallel planes, which does not correspond to the data. Instead, the fit describes the distribution of the TFs as a single layer separated only due to the course of the synaptonemal complex. Yellow: point representation of the transverse filaments, gray: two fitted planes. Point coordinates in pixel.

## Discussion

The aim of this study was to generate detailed quantitative and topological information about the SC structure in the mouse. The availability of higher computational power and the improved sample preparation enabled us to generate tomograms with a high lateral (< 1nm) and z-resolution (3-4 nm) at near-native structure preservation compared to previous SC ET studies. This resolution allowed us, by manual annotation, to create a precise 3D model of the synaptonemal complex. Based on this model, we obtained quantitative data on the individual components of the SC and generated a protein localization map with a special focus on the TFs. Additionally, this novel 3D model enabled us to validate some and challenge other assumptions about the topology of the SC.

The main features and dimensions of the synaptonemal complex are evolutionarily conserved. Strikingly, the width of the central region is approximately 100 nm throughout the animal and plant kingdoms [3]. We measured a CR-width of 114 nm (± 17 nm) which matches the width previously determined and validates the precision of our approach. In addition, the distance we measured from the inner edge of the LE to the inner edge of the CE, as well as the width of the CE that we determined is in accordance with previous EM studies.

While the overall dimensions of the SC have been characterized extensively in yeast, plants, and animals, little quantitative data on the TFs exists. Previously, TF widths ranging from 2 nm to 10 nm, with an average width of 5 nm have been reported. Some of the thicker filaments appeared to split into fibers as thin as 2 nm [39, 38, 35, 36]. Here, we were able to resolve the diameter of TFs with very little deviation (1.9 nm ± 0.3 nm) due to the advanced sample preparation, in combination with the higher resolution that we achieve in 3D through the acquisition of fine-grained double tilt (1° steps) series. This value is in accordance with both the width of a SYCP1 dimer and the lower limit of TF width of previous studies. However, we did not observe the previously reported splitting of thicker TFs into thinner fibers.

Available evidence indicates that two SYCP1 molecules form a long coiled-coil structure of about 900 Å flanked by unstructured N- and C-termini [21]. The C-termini are embedded in the LE, while the N-termini reside within the CE [16, 20]. Here, for the first time, we were able to trace and model the entire course of the 88 nm (±14 nm) long TFs. This value closely matches the length of the SYCP1 coiled-coil structure (see above). Our measurements revealed that significant parts of the TFs, and correspondingly of the coiled-coil of the SYCP1 dimer, are embedded in the lateral and central element, respectively.

Earlier ET studies supported a model of TFs breaching the central region (CR) from one LE to the other. Yet, the authors reported that a subset of filaments only spans half of the central region, ending in the CE [35, 36, 4]. These shorter TFs are in accordance with the localization of the SYCP1 C-terminus to the LE and the SYCP1 N-terminus to the CE. Yet, to resemble the previously reported breaching of TFs, two TFs had to be directly opposing each other, which we do not observe frequently under our experimental conditions. All examined TFs end in the CE with an average minimum distance of 17 nm (± 9 nm) for opposite TFs.

We further determined the number of TFs along a SC as 79 TFs/μm (± 8), which is in accordance with the upper limit of 50-80 TFs/μm previously reported in hamster [38]. As mentioned above, the TFs consist of SYCP1 dimers. Consequently, one micrometer of SC length contains 142 to 174 SYCP1 molecules on average. Interestingly, upon assigning the TFs to the closest LE, we measured a difference in the distribution of up to 21 % comparing the two sides of the central region. This asymmetry implies that a certain TF may lack a partner on the opposite side of the central region. Overall, we observed three types of ‘pairings’ in the CE: the aforementioned opposing pairs, pairs of parallel TFs and TFs apparently lacking a partner (single TFs). The unstructured N-termini (120 amino acids long) of the SYCP1 molecules that mediate the contact between TFs in the CE, cannot be visualized by ET.

Very recently, Dunce and colleagues have proposed a model for TF organization in the central region of the SC. According to their model, the coiled-coil domain of a SYCP1 dimer interacts over a short distance with the coiled-coil of another SYCP1 dimer of the same side, thereby forming a tetrameric structure close to the CE [21]. However, our results do not support this model, as the average minimum distance we measured between parallel TFs is 20 nm (± 10 nm). We further did not detect any branching of the filaments as suggested in the model of Dunce et al.. The differences between the crystallographic and ET-derived models could be due to differences in the experimental approaches. Dunce et al. have investigated the in vitro properties of SYCP1 molecules (i.e. in the absence of other SC proteins) while we have investigated the 3D organization of SYCP1 within the context of the SC.

Within the LE, we determined an average minimum distance of 21 nm (± 10 nm) between two neighboring TFs. Interestingly, this spacing is in accordance with the 20 nm length of the tetrameric core of SYCP3 as resolved by crystallography [13]. Repeating units of the protein assemble into a lattice with N-terminal regions of DNA binding domains exposed at both sides. Biochemical assays confirmed the binding of dsDNA to the N-terminus of SYCP3. The authors integrated these findings into a model that understands SYCP3 as a molecular spacer, which organizes the DNA of the meiotic chromosomes into loops separated by 20 nm [13]. In 2018, Dunce et al. included the assembly of SYCP1 to this model based on their crystallographic and biochemical data. In the mature SC, they propose that the C-terminal ends of all neighboring SYCP1 proteins interact in a U-shape, which coats one loop of DNA per pairing [21]. This assembly suggests an even distribution of TFs along the SC. However, this is not the case according to our data on the distribution of TFs (see also below). We propose a model where one SYCP1 dimer matches one tetramer of SYCP3. In this scenario, SYCP3 would act as both the previously suggested molecular spacer for chromatin loop organization as well as a spacer for possible insertion slots for SYCP1 into the lateral element. While we suggest that SYCP3 could dictate the minimum distance between TFs in the LE, it is noteworthy that no direct interaction between SYCP1 and SYCP3 has been shown [17, 11, 16]. SYCP2 as the other major component of the LE interacts both with SYCP1 and SYCP3 and could convey the spacing in SYCP1 dimer and thereby TF insertion slots predefined by repetitive SYCP3 units [18, 40]. Nevertheless, additional factors might be involved in TF distribution as suggested by the analysis of *Syce3*^−/−^ mice. Meiocytes can assemble SC-like structures between homologous chromosomes that lack LEs, but show TFs and a CE [41].

Previous immunoelectron and immunofluorescence studies provided evidence for a bilayered organization of TFs in mouse SCs. Visual inspection and mathematical modeling of our ET-derived 3D models did not confirm this view. The reasons for this discrepancy may be due to the different experimental approaches used. Here, we have directly visualized and analyzed the topology of hundreds of individual TFs. In the previous studies, TF epitopes were localized by means of antibodies and indirect immunolocalization methods [i.e. 33, 31]. As discussed by Schücker et al., restricted epitope accessibility might be a caveat [33]. In the case of EM post-embedding methods [31], major problems are epitope density and the fact that antibodies do not penetrate the plastic sections. As already discussed, a robust workflow is necessary to use immuno-EM as a quantitative tool [42].

The work described here provides the first quantitative 3D model of the TF assembly in the framework of the SC. Major unexpected outcomes were the asymmetric distribution of TFs and the absence of layers. Here, we propose that these features might be related to the dynamic properties of the SC. SCs are helical structures that move and bend in the nuclear space due to telomere movements at the plane of the nuclear envelope. We envision that the TF distribution described here (i.e. opposing a rigid symmetry in TF order) would favor the flexibility of the SC structure while improving resistance to mechanical stress. This implies that during meiotic prophase TFs might be dynamic in the sense of local assembly/disassembly and distribution in response to and/or allowing SC dynamics. Certainly, further studies are required to understand the dynamic properties of TFs.

## Methods

### Ethics Statement

Animal care and experiments were conducted in accordance with the guidelines provided by the German Animal Welfare Act (German Ministry of Agriculture, Health and Economic Cooperation). Animal housing and breeding was approved by the regulatory agency of the city of Würzburg (Reference ABD/OA/Tr; according to §11/1 No. 1 of the German Animal Welfare Act).

### Testis tissue preparation for electron microscope tomography

14-day-old male C57.6J/Bl6 wildtype were sacrificed using CO_2_, followed by cervical dislocation. We then extracted the seminiferous tubules, pre-fixed the tissue with Karnovsky fixative [43] and additionally cryo-immobilized it through high-pressure freezing. The protocol was adapted from an approach published by Dhanyasi et al. [44]. For the high-pressure freezing, we used 10% BSA in PBS as a filler/ cryo-protectant and freezing platelets of 200 μm (carrier type A with a 0 μm recess carrier type B as a lid) in depth. Freezing speed was >20,000 K/s, the pressure > 2100 bar in an EM HPM100 (Leica, Wetzlar, Germany). The tissue was freeze substituted according to previously published protocols using a Leica EM AFS2 freeze substitution system (Leica, Wetzlar, Germany; for freeze substitution protocol refer to supporting information) [45, 46]. We embedded the tubules in epoxy resin and cut tissue sections of 250 nm thickness with a Histo Diamond Knife (Diatome, Biel, Switzerland). The sections were transferred to copper grids and contrasted with 2.5 % uranyl acetate in Milli-Q or ethanol, followed by 50 % Reynolds’ lead citrate in Milli-Q [47]. A thin layer of carbon was added to the sections to avoid charging during tomogram acquisition. 18 nm colloidal gold coupled to guinea pig IgG was used as an unspecific fiducial marker to aid in tomogram reconstruction.

### Electron tomography (ET)

We acquired dual axis tilt series of the synaptonemal complex in pachytene spermatocytes at a magnification of 40,000x with a JEOL JEM-2100 transmission electron microscope (JEOL, Munich, Germany). The microscope is equipped with a TVIPS TemCam F416 4k×4k camera (Tietz Video and Imaging Processing Systems, Gauting, Germany) and was operated at 200kV. Each tilt axis consists of 141 consecutive images taken in 1° increments from −70° to 70° with SerialEM [48]. Samples were irradiated for 10 minutes prior to acquisition to minimize drift.

### Tomogram generation and data processing

Six tomograms were reconstructed through weighted-back projection with etomo. We segmented the structures of interest (lateral elements, central element and transverse filaments) manually in 3dmod. Both etomo and 3dmod are part of the IMOD software suite [49]. 3dmod contains a measuring tool, which we used to calculate the width of the transverse filaments and the length of the synaptonemal complex. The length and number of the transverse filaments were extracted with the command line extension imodinfo. Another command line program, model2point, was used to generate a point representation of each segmented structure making up the 3D model. All further values were computed on this point representation using a custom Matlab script. Three tomograms of the total six tomograms were acquired of synaptonemal complexes attached to the nuclear envelope and three in the center of the cell to avoid detecting differences in SC structure at a site of greater mechanical stress (SCs at NE) compared to sites of less mechanical stress (interstitial SCs). Initially attached and interstitial SCs were treated as separate data sets. Our measurements did not reveal significant differences between the two sets (data not shown). Consequently, in this study, measurements of all six tomograms were treated as one data set.

For each filament, we determined the intersection point with the corresponding lateral element and the central element. Based on these intersection points we calculated the length of the transverse filaments within the lateral element as the Euclidean distance between the intersection point and the respective endpoint of each filament. Similarly, we determined the length of the transverse filaments in the central region as the Euclidean distance between the intersection with the central element and the intersection with the lateral element. The total length of the transverse filaments was additionally calculated as the Euclidean distance between its endpoints.

In order to determine the width of the central region, we computed for every intersection point in the lateral element the shortest distance to an intersection point of the opposing lateral element. We determined the distance between the central and lateral element as well as the width of the central element accordingly.

We calculated the minimum distance between transverse filaments in the lateral element by determining the Euclidean distances between a filament’s endpoint in the LE and the endpoints of all other transverse filaments. We repeated this process for the endpoints in the central element. We hereby determined the shortest distance between the endpoints on the same side and the shortest distance to an endpoint of a TF of the opposite side.

In order to determine the number of TFs per μm of synaptonemal complex length or per μm^2^ of SC area we first projected the endpoints of the TFs onto a 2D plane. We then fitted a rectangle to the maxima and minima of these points. From the dimension of the rectangle, we determined the average length and area of the synaptonemal complex occupied by TFs.

To check for a multilayer organization of TFs within the synaptonemal complex, we fitted two planes to the two endpoints of the filaments on either side of the central element and visually evaluated for a parallel orientation of the two planes.

## Author contributions

R.B. and M-C.S. designed research; M-C.S. and S.F. performed research; M-C.S. and S.F. analyzed the data; C.S. contributed protocols/facilities; M-C.S. and R.B. wrote the manuscript; M-C.S., S.F, C.S. and R.B. revised the manuscript.

## Author declaration

The authors declare no conflict of interest

## Acknowledgements

We thank Markus Engstler for generous support, Silke Braune and Elisabeth Meyer-Natus for excellent technical assistance, Irene da Cruz for helpful discussions and the members of the Imaging Core Facility for technical advice. Supported by grant Be1168/8-1 of the German Science Foundation to R.B.

